# Swarming rate and timing of unmanaged honeybee colonies (*Apis mellifera carnica*) in a forest environment

**DOI:** 10.1101/2024.09.07.611535

**Authors:** Benjamin Rutschmann, Patrick L. Kohl, Ingolf Steffan-Dewenter

## Abstract

Investigating the life history of social insect colonies and the demography of their populations are important for their conservation, but data collection is challenging. There is a growing interest in understanding the population status of wild-living honeybee colonies across Europe, for which it is critical to collect data on survival and natality rates. Although survival rates can be investigated through regular inspections of wild nests, the accurate quantification of natality rates (i.e., the number of swarms produced per colony per year) remains a significant challenge. Using digital weight scales, we remotely monitored the natural swarming behavior of ten unmanaged *Apis mellifera carnica* colonies housed in static-volume hives (45L) in a forest region of southern Germany. During the 2019 season, between mid-May and late June, we recorded 17 swarming events, averaging 1.7 swarms per colony. Our observations offer a reference point for the timing, frequency, and size of honeybee swarms that helps us understand the natural reproductive patterns of wild-living honeybees in a temperate forest environment.

## INTRODUCTION

Understanding the life history strategy of species and studying the demography of their populations are core tasks in conservation biology. For an analysis of the viability of a given population, two variables are key: survival and natality. Survival (or its converse, mortality) refers to the proportion of the population that endures from one year to the next, influenced by factors such as predation, diseases, and other environmental conditions (e.g. weather conditions or resource availability). Conversely, natality encompasses the introduction of new individuals into the population through reproduction, contributing to population growth (Krebs, 1972).

Although European populations of the western honeybee (*Apis mellifera*) have traditionally been viewed through the lens of apiculture, recent studies have highlighted the existence of wild-living^1^ honeybees across various regions (e.g. Oleksa et al., 2013; Kohl and Rutschmann, 2018; Requier et al., 2020; Browne et al., 2020; Bila Dubaić et al., 2021; Rutschmann et al., 2022; Visick and Ratnieks, 2023; Rutschmann et al., 2024). Understanding the population demography of these wild-living colonies is crucial for comprehending their ecological and evolutionary roles (Kohl and Rutschmann, 2018; Kohl et al., 2022). Furthermore, if self-sustaining populations were discovered, these would offer unique insights into how honeybees can persist despite challenges like parasitic pressures which, in turn, could contribute to the development of more sustainable beekeeping practices (Panziera et al., 2022; Requier et al., 2019).

Honeybee colonies are intrinsically long-lived (multiple years) since young queens can supersede old queens as the reproductive individual of the society. In temperate regions, honeybees exhibit high site fidelity, allowing survival rates to be assessed through repeated visits to their nest sites (Seeley, 1978; Oldroyd et al., 1997; Rutschmann et al., 2022; Kohl et al., 2022; Lang et al., 2022; Cordillot, 2024; Rutschmann et al., 2024). Acquiring comprehensive and reliable data on the annual natality rates of honeybee colonies within a population, however, poses a significant challenge (Otis and Wearing-Wilde, 1992). Colony reproduction is achieved through swarming, a process in which a single colony divides into two or more distinct colonies under favorable environmental conditions. In preparation of this process, the colony raises several young queens. Before these new queens emerge, the old queen leaves with more than 70% of the colony’s worker bees as the so-called prime swarm (Seeley, 1985; Winston, 1991). Subsequently, several afterswarms–each with at least one of the newly emerged young queens and another portion of the workers–may leave the original nest site to found independent colonies (Otis and Wearing-Wilde, 1992; Seeley, 1985; Winston, 1991). The annual natality rate in honeybees is thus defined as the frequency of swarms (prime swarm plus afterswarms) per colony per year.

Although a queen change during the swarming season (which is typically indicative of a swarming event) can be detected using genetic maternity tests of workers, or, in case of colonies managed in hives by directly monitoring the presence of color-marked queens, these methods fall short in accurately determining the number of afterswarms (Otis and Wearing-Wilde, 1992). Therefore, prior studies on annual swarming rates required continuous observation of nest entrances (Otis and Wearing-Wilde, 1992), a method that is notably time-consuming.

We here demonstrate the use of digital weight scales to remotely and accurately monitor honeybee colony swarming events. We gathered data from ten colonies that were kept in hives with a static volume of approximately 45 liters in a forest region in Germany. By avoiding beekeeping manipulations, we ensured that our observations reflected near natural behavior. The method not only enhances previous research techniques but also adds to our understanding of the natural timing and frequency of swarming and, consequently, the population dynamics of wild-living honeybee populations in a European forest environment.

## METHODS

### Study area

This study was conducted in the Steigerwald, a low mountain forest region in southern Germany. The details of our experimental setup are described in Rutschmann et al. (2023). The managed forested area covers approximately 165 km^2^ and primarily consists of beech trees (*Fagus sylvatica*, 44%), followed by oak (*Quercus* spp., 21%), Scots pine (*Pinus sylvestris*, 13%), spruce (*Picea abies*), and hornbeam (*Carpinus betulus*) (Mergner and Kraus, 2020). Surrounding grassland areas feature many orchard meadows, and the most prevalent crops include cereals, corn, and oilseed rape.

### Honeybee colonies, weight scales and statistical analysis

In July 2018, we selected 12 equally sized *Apis mellifera carnica* colonies, each featuring a one-year-old queen. The colonies were housed in single 10-frame Zander hive boxes whose volume of approximately 45 liters matches the natural cavity volume preferences of temperate *Apis mellifera* (Seeley and Morse, 1976). We placed the colonies at various sites inside the forest or along its edges. Spatial independence was achieved by placing neighboring colonies around 2 km or more apart, as detailed in Rutschmann et al. (2023). Each hives was set up on a weight scale (Capaz BEE HIVE SCALE GSM 200) that recorded hive weight once every hour and forwarded data via mobile communication.

The only beekeeping interventions conducted included treatments against varroosis using formic acid in August 2018 (one year prior to the swarming season) and the temporary placement of pollen traps for 1–3 days each month during the foraging season. During preliminary data analysis, we addressed potential weight artifacts arising from hive maintenance manipulations. Due to the malfunctioning of weight scales at two locations, our investigation was limited to recordings from 10 of the 12 initially selected colonies. Swarm departures were identified as abrupt weight losses exceeding 500 grams within an hour. We calculated swarm weight (as a measure of swarm size) by comparing hive weights the hour before and after the swarm’s departure. All swarms issued throughout the 2019 season were recorded. To evaluate the weather conditions during swarming events, we retrieved average daily temperatures and precipitation data from nearby weather stations, provided by Agrarmeteorologie Bayern (www.wetter-by.de).

To assess the influence of swarm category (prime or afterswarm) on swarm weight, we performed a linear mixed-effects model analysis using the lme4 package in R (Bates et al., 2015). Weight was the response variable, swarm category was the predictor, and colony ID was included as a random effect to account for variability between different colonies. Additional models incorporating Julian date were tested but did not improve the explanatory power of the model (Figure SI1). Residuals of the models were inspected with DHARMa package (Hartig and Hartig, 2017). For inference, a Type II ANOVA was performed using the Anova function from the car package (Fox et al., 2012). All statistical analyses were performed using R software (version 4.3.1; R Core Team, 2016). For data processing and graphical representation of the results, we utilized ‘tidyverse’, ‘ggplot2’, ‘patchwork’ and ‘ggpol’ (Wickham, 2017, 2016; Pedersen, 2020; Tiedemann, 2020).

## RESULTS

We recorded a total of 17 swarming events across ten colonies of *Apis mellifera carnica* during the 2019 swarming season, resulting in an average of 1.7 (SD: ±0.8) swarms per colony (1 prime swarm and 0.7 afterswarms) (Figure 1A). All colonies swarmed at least once: five colonies issued a single swarm; three colonies produced two swarms (a prime swarm and one afterswarm); and two colonies gave rise to three swarms (a prime swarm and two afterswarms). Swarms departed between May 17 and June 27, and the median swarming date was May 30 (Figure 1B). The interval between prime swarms and afterswarms averaged 9 ± 3 days (range: 6–13 days, N=5). For the two colonies that produced two afterswarms, the interval between the first and second afterswarm were 4 and 8 days. The median departure time for swarms was 11:00 AM (range: 8:30 AM to 5:30 PM). Swarming usually occurred when it was getting warmer, following a period of relative cold, and importantly, on days without rain (Figure 2).

**Figure 1:**
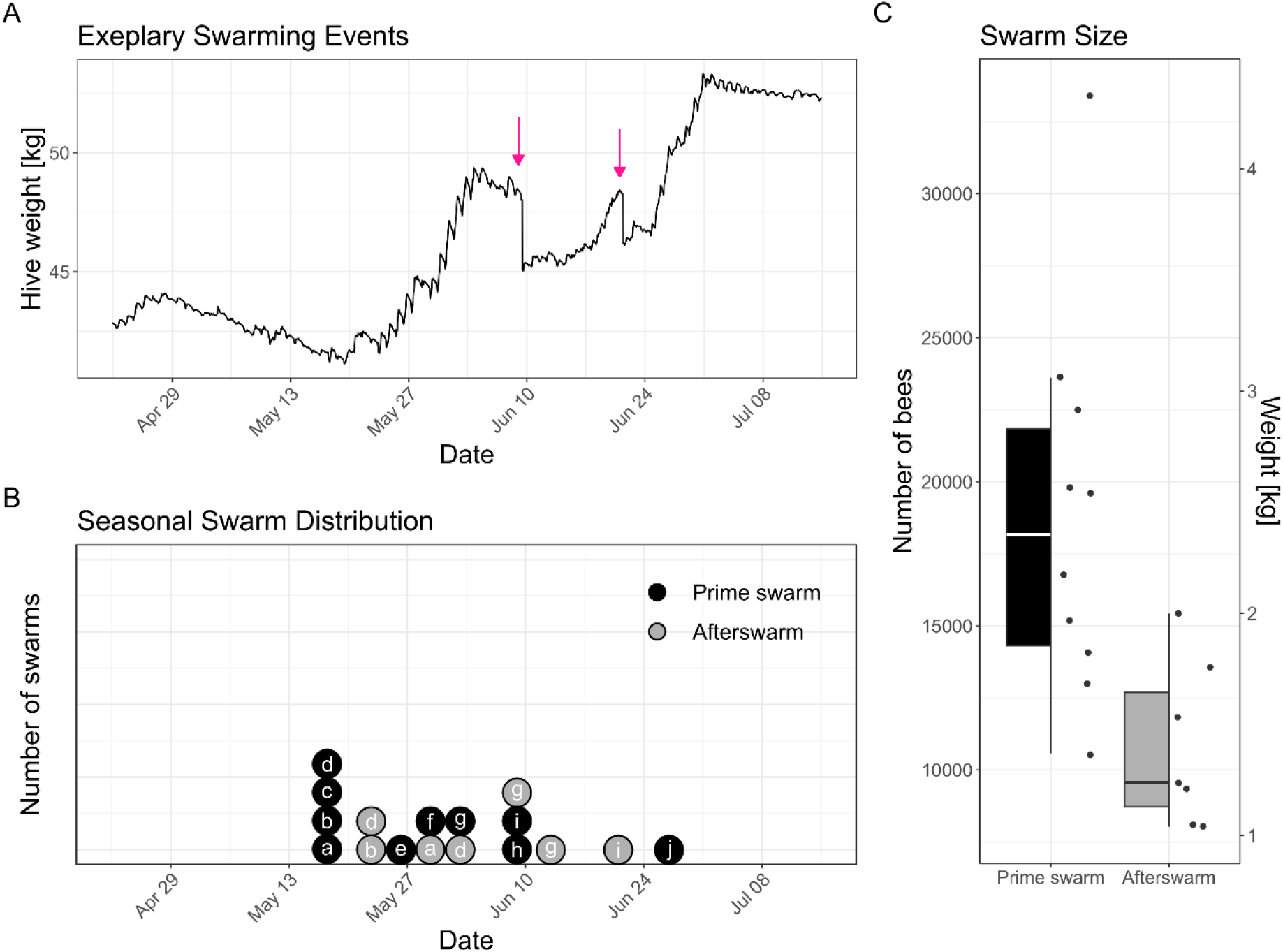
Overview of swarming activity in a forest environment and its characteristics. A) Provides an example of hive weight recordings from colony “i” during the swarming period, where pink arrows indicate the occurrence of a prime swarm and an afterswarm. B) Shows the temporal distribution of swarms throughout the season, with letters representing the different parental colonies. C) A boxjitter plot displays the size and weight of the recorded prime swarms and afterswarms.

**Figure 2:**
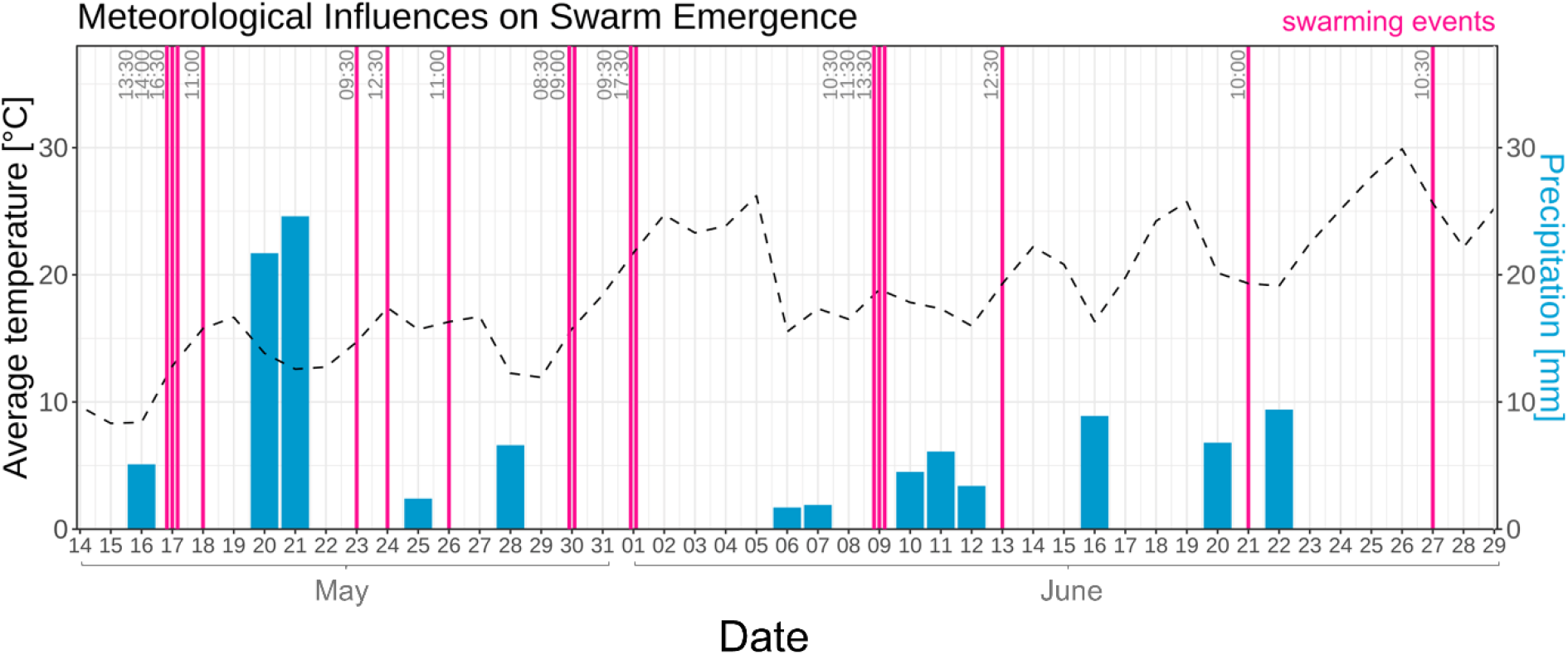
Average temperature and precipitation in the study area from mid-May to late June 2019. The blue bars represent daily precipitation in millimeters, while the black dashed line represents the daily average temperature in degrees Celsius. Pink vertical lines indicate the occurrence of swarming events, with the corresponding time of day labelled.

We observed notable variation in swarm weight, with an overall mean and SD weight of 2.01 ± 0.86 kg (range: 1.04 kg – 4.33 kg). This variability stemmed mainly from the significant differences between prime swarms, averaging 2.44 ± 0.86 kg, and afterswarms, at 1.40 ± 0.37 kg (Figure 1C). Utilizing Mitchell (1970) empirical formula (7,716 bees = 1 kg), we estimated an average of 18,843 bees in prime swarms and 10,836 bees in afterswarms, highlighting the statistically significant variation in swarm sizes (χ^2^ = 21.276, df = 1, p < 0.001).

## DISCUSSION

Our study provides insights into the natural swarming behavior and the natality rates of unmanaged *Apis mellifera carnica* colonies in a German forest environment. It offers a critical reference point for ongoing and future demographic investigations of wild-living honeybees in Europe.

### Comparison with other studies

The observed average of 1.7 swarms falls within the overall range of 0–4 swarms per colony and year recorded for temperate honeybees (reviewed by Winston, 1991). However, a comparison between individual studies reveals notable variations between swarming rates.

This highlights the impact of geographical location and subspecies differences on swarming frequency (Uzunov et al., 2014).

In Kansas, USA, Winston (1980) recorded 3.6 daughter colonies per year from a combination of *Apis mellifera carnica* and *Apis mellifera ligustica* colonies, thus a significantly higher natality rate than reported here. In British Columbia, Canada, Lee and Winston, (1987) observed an annual reproductive rate of 2.2 swarms. Seeley (1978, 2017) reported that the probability for colonies to enter a swarming cycle was around 0.9 in the Arnot forest in New York State. Since this number does not account for afterswarms, it resembles our observations more closely. The lowest propensity to swarm was reported by Fries et al. (2006) for colonies on Gotland, an island in the Baltic Sea, where the proportion of reproducing colonies (with at least one swarm) varied between 0 and 60% in six observation years. Similarly, Le Conte et al. (2007) documented variable swarming rates in untreated colonies resistant to *Varroa destructor* in France, with an average rate of 41.50 ± 9.94% across six years.

Otis and Wearing-Wilde (1992) pointed to the significant contribution of afterswarms to the overall number of swarms produced in a population, and our finding of an average of 0.7 afterswarms per colony supports that conclusion. Five out of ten colonies in our study produced more than the prime swarm. This shows that it is important to use methods that can detect afterswarms to accurately estimate natality rates.

In non-temperate regions, natality rates for other subspecies have been found to be substantially higher (Winston et al. 1983), which underscores the significant role local environmental conditions and biological factors play in shaping reproductive strategies of honeybees (Ruttner, 1988; Nuru et al., 2002; Strange et al., 2007; Norrström et al., 2021). Additionally, factors such as the queen’s age (younger queens exhibit increased pheromone production, thereby more effectively suppressing the development of new queens) and the size of the hive have been demonstrated to influence swarming behavior (Uzunov et al., 2014; Smith et al., 2017). For example Loftus et al. (2016) showed a clear relationship between hive size and swarming propensity; small hives (42 liters) had a higher swarming rate, with 10 out of 12 colonies swarming, compared to larger hives (168 liters), where only 2 out of 12 colonies swarmed.

Besides providing data on swarming rate, this study points to the large variation in swarm sizes that exist within a single season and population, noting marked differences between primary swarms and afterswarms. Prior research revealed a broad spectrum of swarm sizes, ranging from as few as 2,400 bees (0.31 kg) to as many as 41,500 bees (5.33 kg), with mean populations reported at 11,800 bees (1.53 kg) by Fell et al. (1977) and 14,000 bees (1.8 kg) by Burgett and Morse (1974). It must be noted that the smaller worker populations and unmated queens associated with afterswarms lower their survival prospects (Lee and Winston, 1987).

### Swarming and net reproductive rate

Assessing the self-sustainability of a honeybee population necessitates a thorough understanding of its annual survival and natality rates. The net reproductive rate (*R*_0_), defined for honeybees by the formula *R*_0_ *= s + s × n* (Kohl et al., 2022), integrates survival rate (*s*) and natality rate (*n*) (i.e. the number of swarms). This index helps in identifying whether populations are diminishing (*R*_0_ < 1), maintaining stability (*R*_0_ = 1), or experiencing growth (*R*_0_ > 1), intrinsically—i.e., without considering immigration from external sources such as managed apiaries.

If we assume that our finding of 1.7 swarms per colony and year, which is based on a single swarming season and on a small number of colonies, represents well the average reproductive rate in our study system, wild-living honeybee colonies in managed beech-dominated forests in Germany would necessitate a minimum annual survival rate of 37% to achieve *R*_0_ = 1 and thus, population sustainability. Such a survival rate is considerably higher than the observed survival of 10.6% as reported by Kohl et al. (2022) or 12% by Rutschmann et al. (2024). This discrepancy highlights the urgent need for deeper investigations into the factors influencing survival and reproduction in these cohorts, to develop strategies for their conservation and management.

### The role of landscape in the timing of swarming

Resource availability on the landscape scale plays a pivotal role in the survival of wild-living honeybee colonies (Rutschmann et al., 2022; Kohl et al., 2023) and the establishment of new swarms (Seeley, 1978; Seeley and Visscher, 1985; Otis and Wearing-Wilde, 1992; Seeley, 2017). A comparison with two other phenological data sets also suggests that the timing of swarming is affected by landscape context. Henneken et al. (2012), utilizing a crowdsourced dataset of 1,335 swarming events in Germany during 2011, found that swarming began in April with a first peak in early May. In addition, a predominantly urban dataset from 2019, the same year as our study, which included 77 swarm observations (55 from Munich and 22 from across Germany), reported swarming as early as 17 April (Rutschmann et al., 2024).

While the median swarming date in the urban setting was around 19 May 2019 (Remter, Roth, Rutschmann, personal communication), the median swarming date in this study was 30 May 2019. The delayed timing of swarms in our study region might be explained by inferior foraging conditions in forest-dominated compared to agro-urban landscapes. We know from a study on foraging behavior in the same study region that, especially in April and May during an important phase of colony weight gain, honeybees find superior foraging opportunities in open areas such as grassland and cropland (Rutschmann et al. (2023). Colonies nesting in forests might also face cooler temperatures and fewer hours of sunlight compared to colonies in non-forest landscapes and thus might show delayed development in spring due to fewer hours available for foraging (Czekońska et al., 2023). An additional possible influence on the delayed onset of swarming observed in forest settings could be the lack of stimulative feeding practices that are commonly used in urban/agricultural settings by many beekeepers.

Additionally, the temporary placement of pollen traps for 1–3 days each month during the foraging season as part of a different study, may also have contributed to a slight delay.

### Future directions and methodological advancements

The adoption of weight scales for the surveillance of swarming events introduces a non-invasive, efficient approach to studying the reproductive patterns of honeybees. We encourage the broader application of this technology to detect swarming in weight monitoring initiatives such as McMinn-Sauder et al. (2024), Johannesen et al. (2022), Lecocq et al. (2015), Kuchling et al. (2018), Komasilova et al. (2023) or Prešern et al. (2019). While our study focused on a limited number of colonies (N = 10), future research with a larger sample size could investigate the differences in swarming across various landscapes and subspecies, providing deeper insights into the factors influencing honeybee reproduction.

## ACKNLOWEDGMENTS

We express our gratitude to Ulrich Mergner and the Ebrach state forest department for granting us permission to conduct our research in the Steigerwald forest. Our thanks also go to Jürgen Tautz for his generosity in supplying beekeeping materials and additional resources. Special thanks to Gaby Läbisch for her assistance in managing the study colonies before the experimental phase. We are thankful to Susanne Schiele and Gemma Villagomez for their assistance with the queen bees and the weight scales. We acknowledge the environmental impact of our research, specifically the carbon emissions from traveling to the field sites. Hence, the corresponding author has privately compensated the estimated two tons of CO2 emissions.

## DATA AVAILABILITY STATEMENT

Data used in the submitted manuscript can be made available to a limited degree after a reasonable request to the corresponding author.

## AUTHOR CONTRIBUTIONS

BR conceived the study, analysed and visualized the data and wrote the first draft of the manuscript. PLK provided assistance in the field, and ISD provided resources. All authors reviewed and edited the manuscript.

## ETHICS APPROVAL

No honeybees were harmed during this study, which used non-invasive monitoring methods. Ethical approval was not required as honeybees are invertebrates.

## Footnotes

1 To describe honeybee colonies that are not managed and chose their nest site on their own, we use the term “wild-living”.

## REFERENCES

Bates, D., Mächler, M., Bolker, B., Walker, S., 2015. Fitting Linear Mixed-Effects Models Using lme4. Journal of Statistical Software 67, 1–48. 10.18637/jss.v067.i01

Bila Dubaić, J., Simonović, S., Plećaš, M., Stanisavljević, L., Davidović, S., Tanasković, M., ćetković, A., 2021. Unprecedented Density and Persistence of Feral Honey Bees in Urban Environments of a Large SE-European City (Belgrade, Serbia). Insects 12, 1127. 10.3390/insects12121127

Browne, K.A., Hassett, J., Geary, M., Moore, E., Henriques, D., Soland-Reckeweg, G., Ferrari, R., Mac Loughlin, E., O’Brien, E., O’Driscoll, S., Young, P., Pinto, M.A., McCormack, G.P., 2020. Investigation of free-living honey bee colonies in Ireland. Journal of Apicultural Research 1–12. 10.1080/00218839.2020.1837530

Burgett, D.M., Morse, R.A., 1974. The Time of Natural Swarming in Honey Bees1, 2. Annals of the Entomological Society of America 67, 719–720. 10.1093/aesa/67.4.719

Cordillot, F., 2024. Erste Suche nach wilden Honigbienen (Apis mellifera L., 1758) auf der Schweizer Alpennordseite. Entomo Helvetica 17, 97–114.

Czekońska, K., Łopuch, S., Miścicki, S., 2023. The effect of meteorological and environmental variables on food collection by honey bees (Apis mellifera). Ecological Indicators 156, 111140. 10.1016/j.ecolind.2023.111140

Fell, R.D., Ambrose, J.T., Burgett, D.M., De Jong, D., Morse, R.A., Seeley, T.D., 1977. The Seasonal Cycle of Swarming in Honeybees. Journal of Apicultural Research 16, 170–173. 10.1080/00218839.1977.11099883

Fox, J., Weisberg, S., Adler, D., Bates, D., Baud-Bovy, G., Ellison, S., Firth, D., Friendly, M., Gorjanc, G., Graves, S., others, 2012. Package ‘car.’ Vienna: R Foundation for Statistical Computing 16.

Fries, I., Imdorf, A., Rosenkranz, P., 2006. Survival of mite infested (Varroa destructor) honey bee (Apis mellifera) colonies in a Nordic climate. Apidologie 37, 564–570. 10.1051/apido:2006031

Hartig, F., Hartig, M.F., 2017. Package ‘DHARMa.’

Henneken, R., Helm, S., Menzel, A., 2012. Meteorological Influences on Swarm Emergence in Honey Bees (Hymenoptera: Apidae) as Detected by Crowdsourcing. env. entom. 41, 1462–1465. 10.1603/EN12139

Johannesen, J., Wöhl, S., Berg, S., Otten, C., 2022. Annual Fluctuations in Winter Colony Losses of Apis mellifera L. Are Predicted by Honey Flow Dynamics of the Preceding Year. Insects 13, 829. 10.3390/insects13090829

Kohl, P.L., Rutschmann, B., 2018. The neglected bee trees: European beech forests as a home for feral honey bee colonies. PeerJ 6, e4602. 10.7717/peerj.4602

Kohl, P.L., Rutschmann, B., Sikora, L.G., Wimmer, N., Zahner, V., D’Alvise, P., Hasselmann, M., Steffan-Dewenter, I., 2023. Parasites, depredators, and limited resources as potential drivers of winter mortality of feral honeybee colonies in German forests. Oecologia 202, 465–480. 10.1007/s00442-023-05399-6

Kohl, P.L., Rutschmann, B., Steffan-Dewenter, I., 2022. Population demography of feral honeybee colonies in central European forests. R. Soc. open sci. 9, 220565. 10.1098/rsos.220565

Komasilova, O., Zacepins, A., Kviesis, A., Komasilovs, V., Ozols, N., 2023. Comparing weight dynamics between urban and rural honey bee colonies in Latvia 556. 9Kb. 10.15159/AR.23.019

Krebs, C.J., 1972. Ecology. The experimental analysis of distribution and abundance.

Kuchling, S., Kopacka, I., Kalcher-Sommersguter, E., Schwarz, M., Crailsheim, K., Brodschneider, R., 2018. Investigating the role of landscape composition on honey bee colony winter mortality: A long-term analysis. Sci Rep 8, 12263. 10.1038/s41598-018-30891-y

Lang, U., Albouy, V., Zewen, C., 2022. Comparative monitoring of free-living honey bee colonies in three Western European regions. Natural Bee Husbandry Magazine.

Le Conte, Y., de Vaublanc, G., Crauser, D., Jeanne, F., Rousselle, J.-C., Bécard, J.-M., 2007. Honey bee colonies that have survived Varroa destructor. Apidologie 38, 566–572. 10.1051/apido:2007040

Lecocq, A., Kryger, P., Vejsnæs, F., Bruun Jensen, A., 2015. Weight Watching and the Effect of Landscape on Honeybee Colony Productivity: Investigating the Value of Colony Weight Monitoring for the Beekeeping Industry. PLOS ONE 10, e0132473. 10.1371/journal.pone.0132473

Lee, P.C., Winston, M.L., 1987. Effects of reproductive timing and colony size on the survival, offspring colony size and drone production in the honey bee (Apis mellifera). Ecological Entomology 12, 187–195. 10.1111/j.1365-2311.1987.tb00997.x

Loftus, J.C., Smith, M.L., Seeley, T.D., 2016. How Honey Bee Colonies Survive in the Wild: Testing the Importance of Small Nests and Frequent Swarming. PLOS ONE 11, e0150362. 10/f8vmd5

McMinn-Sauder, H.B.G., Colin, T., Gaines Day, H.R., Quinlan, G., Smart, A., Meikle, W.G., Johnson, R.M., Sponsler, D.B., 2024. Next-generation colony weight monitoring: a review and prospectus. Apidologie 55, 13. 10.1007/s13592-023-01050-8

Mergner, U., Kraus, D., 2020. Learning from nature: Integrative forest management in Ebrach, Germany. pp. 196–213.

Mitchell, C., 1970. Weights of workers and drones. Amer Bee J.

Norrström, N., Niklasson, M., Leidenberger, S., 2021. Winter weight loss of different subspecies of honey bee Apis mellifera colonies (Linnaeus, 1758) in southwestern Sweden. PLoS ONE 16, e0258398. 10.1371/journal.pone.0258398

Nuru, A., Amssalu, B., Hepburn, H.R., Radloff, S.E., 2002. Swarming and migration in the honey bees (Apis mellifera) of Ethiopia. Journal of Apicultural Research 41, 35–41. 10.1080/00218839.2002.11101066

Oldroyd, B.P., Thexton, E.G., Lawler, S.H., Crozier, R.H., 1997. Population demography of Australian feral bees (Apis mellifera). Oecologia 111, 381–387. 10.1007/s004420050249

Oleksa, A., Gawroński, R., Tofilski, A., 2013. Rural avenues as a refuge for feral honey bee population. Journal of Insect Conservation 17, 465–472. 10/f4xf44

Otis, G.W., Wearing-Wilde, J.M., 1992. Net reproductive rate of unmanaged honeybee colonies, (Apis mellifera L.). Ins. Soc 39, 157–165. 10.1007/BF01249291

Panziera, D., Requier, F., Chantawannakul, P., Pirk, C.W.W., Blacquière, T., 2022. The Diversity Decline in Wild and Managed Honey Bee Populations Urges for an Integrated Conservation Approach. Front. Ecol. Evol. 10, 767950. 10.3389/fevo.2022.767950

Pedersen, T.L., 2020. patchwork: The Composer of Plots.

Prešern, J., Mihelič, J., Kobal, M., 2019. Growing stock of nectar- and honeydew-producing tree species determines the beekeepers’ profit. Forest Ecology and Management 448, 490–498. 10.1016/j.foreco.2019.06.031

R Core Team, 2016. R: A Language and Environment for Statistical Computing. R Foundation for Statistical Computing, Vienna, Austria.

Requier, F., Garnery, L., Kohl, P.L., Njovu, H.K., Pirk, C.W.W., Crewe, R.M., Steffan-Dewenter, I., 2019. The Conservation of Native Honey Bees Is Crucial. Trends in Ecology & Evolution 34, 789–798. 10.1016/j.tree.2019.04.008

Requier, F., Paillet, Y., Laroche, F., Rutschmann, B., Zhang, J., Lombardi, F., Svoboda, M., Steffan-Dewenter, I., 2020. Contribution of European forests to safeguard wild honeybee populations. CONSERVATION LETTERS 13. 10.1111/conl.12693

Rutschmann, B., Kohl, P.L., Machado, A., Steffan-Dewenter, I., 2022. Semi-natural habitats promote winter survival of wild-living honeybees in an agricultural landscape. Biological Conservation 266, 109450. 10.1016/j.biocon.2022.109450

Rutschmann, B., Kohl, P.L., Steffan-Dewenter, I., 2023. Foraging distances, habitat preferences and seasonal colony performance of honeybees in Central European forest landscapes. Journal of Applied Ecology 60, 1056–1066. 10.1111/1365-2664.14389

Rutschmann, B., Remter, F., Roth, S., 2024. Assessing nest sites, survival rates and population dynamics of free-living honeybee colonies in Germany: A comparative study using personally collected and Citizen Science data. bioRxiv. 10.1101/2024.08.02.606354

Ruttner, F., 1988. Biogeography and Taxonomy of Honeybees. Springer Berlin Heidelberg.

Seeley, T.D., 2017. Life-history traits of wild honey bee colonies living in forests around Ithaca, NY, USA. Apidologie 48, 743–754. 10.1007/s13592-017-0519-1

Seeley, T.D., 1985. Honeybee Ecology: A Study of Adaptation in Social Life: A Study of Adaptation in Social Life, Princeton Legacy Library. Princeton University Press.

Seeley, T.D., 1978. Life history strategy of the honey bee, Apis mellifera. Oecologia 32, 109–118. 10/fk89×7

Seeley, T.D., Morse, R.A., 1976. The nest of the honey bee (Apis mellifera L.). Insectes Sociaux 23, 495–512. 10/b76nb3

Seeley, T.D., Visscher, P.K., 1985. Survival of honeybees in cold climates: the critical timing of colony growth and reproduction. Ecol Entomol 10, 81–88. 10.1111/j.1365-2311.1985.tb00537.x

Smith, M.L., Koenig, P.A., Peters, J.M., 2017. The cues of colony size: how honey bees sense that their colony is large enough to begin to invest in reproduction. Journal of Experimental Biology 220, 1597–1605. 10.1242/jeb.150342

Strange, J.P., Garnery, L., Sheppard, W.S., 2007. Persistence of the Landes ecotype of Apis mellifera mellifera in southwest France: confirmation of a locally adaptive annual brood cycle trait. Apidologie 38, 259–267. 10.1051/apido:2007012

Tiedemann, F., 2020. ggpol: Visualizing Social Science Data with “ggplot2.”

Uzunov, A., Costa, C., Panasiuk, B., Meixner, M., Kryger, P., Hatjina, F., Bouga, M., Andonov, S., Bienkowska, M., Conte, Y.L., Wilde, J., Gerula, D., Kiprijanovska, H., Filipi, J., Petrov, P., Ruottinen, L., Pechhacker, H., Berg, S., Dyrba, W., Ivanova, E., Büchler, R., 2014. Swarming, defensive and hygienic behaviour in honey bee colonies of different genetic origin in a pan-European experiment. Journal of Apicultural Research 53, 248–260. 10.3896/IBRA.1.53.2.06

Visick, O.D., Ratnieks, F.L.W., 2023. Ancient, veteran and other listed trees as nest sites for wild-living honey bee, Apis mellifera, colonies. J Insect Conserv. 10.1007/s10841-023-00530-7

Wickham, H., 2017. The tidyverse. R package ver 1, 836.

Wickham, H., 2016. ggplot2: Elegant Graphics for Data Analysis. Springer-Verlag New York.

Winston, M.L., 1991. The biology of the honey bee. harvard university press.

Winston, M.L., 1980. Swarming, afterswarming, and reproductive rate of unmanaged honeybee colonies (Apis mellifera). Ins. Soc 27, 391–398. 10.1007/BF02223731

Winston, M.L., Taylor, O.R., Otis, G.W., 1983. Some Differences between Temperate European and Tropical African and South American Honeybees. Bee World 64, 12–21. 10.1080/0005772X.1983.11097902

