## Supplementary Information for "Swarming rate and timing of unmanaged honeybee colonies (*Apis mellifera carnica*) in a forest environment"

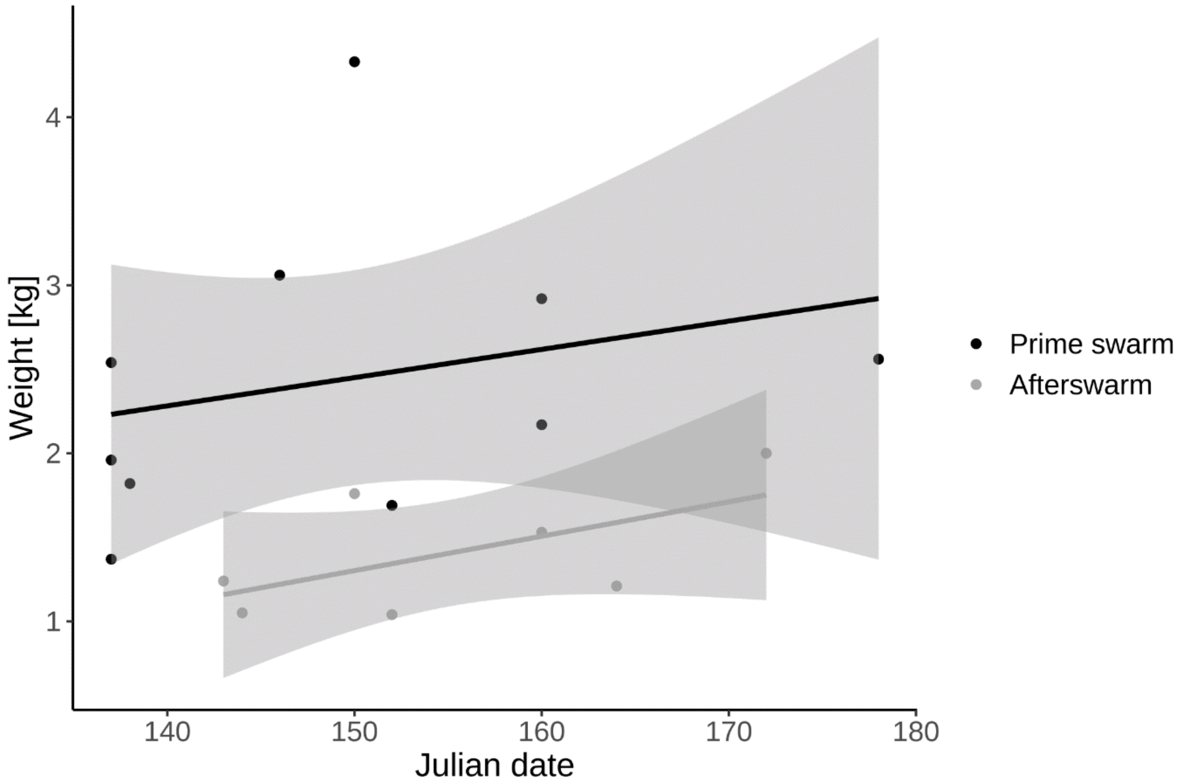


*Figure SI1: Relationship between the weight of swarms and Julian date for prime swarms and afterswarms. The black points and regression line represent prime swarms, while the grey points and regression line represent afterswarms. The shaded areas indicate the standard error (SE) of the regression lines. Including Julian date in the model did not improve the explanatory power of the model.*
